# Experimental system and image analysis software for high throughput phenotyping of mycorrhizal growth response in *Brachypodium distachyon*

**DOI:** 10.1101/779330

**Authors:** Felicia Maviane-Macia, Camille Ribeyre, Luis Buendia, Mégane Gaston, Mehdi Khafif, Fabrice Devoilles, Nemo Peeters, Benoit Lefebvre

## Abstract

- Plant growth response to Arbuscular Mycorrhizal (AM) fungi is variable and depends on genetic and environment factors that still remain largely unknown. Identification of these factors can be envisaged using high-throughput and accurate plant phenotyping.
- We setup experimental conditions based on a two-compartment system allowing to measure *Brachypodium distachyon* mycorhizal growth response (MGR) in an automated phenotyping greenhouse. We developed a new image analysis software *“IPSO Phen”* to estimate of *B. distachyon* aboveground biomass.
- We found a positive MGR in the *B. distachyon* Bd3-1 genotype inoculated with the AM fungi *Rhizophagus irregularis* only if nitrogen and phosphorus were added together in the compartment restricted to AM fungi. Using this condition, we found genetic diversity in *B. distachyon* for MGR ranging from positive to negative MGR depending on the plant genotype tested.
- Our result on the interaction between nitrogen and phosphorus for MGR in *B. distachyon* opens new perspectives about AM functioning. In addition, our open-source software allowing to test and run image analysis parameters on large amount of images generated by automated plant phenotyping facilities, will help to screen large panels of genotypes and environmental conditions to identify the factors controlling the MGR.

## Introduction

Arbuscular mycorrhiza (AM) is one of the most ancient plant-microorganism mutualist symbiosis that occurs between most plant species and Glomeromycota fungi. It is commonly accepted that AM can lead to a better plant growth (Smith & Read, 2008). The extended mycelium network in soil enables the fungi to uptake macro- and micronutrients in areas that are not exploited by plants roots. These nutrients are then transported through the mycelium to the roots and exchanged with plant metabolites (carbohydrates and lipids). Both nitrogen (N) and phosphorus (P), the most limiting nutrients for plant growth can be provided to plants (Pearson & Jakobsen, 1993; Tobar *et al*., 1994; Smith *et al*., 2004; Govindarajulu *et al*., 2005; Fellbaum *et al*., 2014). AM fungi have both nitrate and ammonium transporters and can thus uptake these two N-containing ions from the soil (reviewed in Chen *et al*., 2018). Consequently AM plays a key role in plant nutrition in natural ecosystems and in low fertilization agricultural systems in which the soil N and P availability is often limiting for plant growth.

However, there is host and symbiont genetic variability for plant growth stimulation by AM fungi in controlled conditions. For example, mycorrhizal growth response (MGR) can be either positive or negative in wheat (Hetrick & Wilson, 1992; Xavier & Germida, 1998; Lehnert *et al*., 2018) and sorghum (Watts-Williams *et al*., 2019). Very little is known about the genetic determinants that control MGR. Identification of the underlying plant loci/genes is a major scientific challenge and would help to breed crops for improved MGR and increased performance in low fertilization agriculture.

Genetic studies to identify such loci/genes require high-throughput and accurate plant phenotyping in the presence or absence of the AM fungi. We test the feasibility of such genetic studies on *Brachypodium distachyon*, using an automated phenotyping platform (the “Toulouse Plant-Microbe Phenotyping” or “TPMP” facility).

*B. distachyon* is a wild Mediterranean diploid grass with a relatively small size and short cycle (Girin *et al*., 2014). Variability of *B. distachyon* MGR to various AM fungal genotypes has been shown (Hong *et al*., 2012) but host genetic effect on MGR has not been studied yet.

Effect of AM on plant growth is highly dependent on environmental conditions and in particular on nutrient availability in the soil. Plant colonization by AM fungi is also regulated by nutrient availability (Bonneau *et al*., 2013; Nouri *et al*., 2014). Thus controlling soil nutrient availability is critical to study MGR. We used a two-compartment system, one of them being only accessible to AM fungi and supplemented with N and/or P, to mimic the ability of AM fungi to collect these nutrients in large soil volumes that cannot be explored by the plant root. A compartment only accessible to AM fungi was used to demonstrate N and P transport in plants using ^32^P/^33^P or ^15^N (Pearson & Jakobsen, 1993; Tobar *et al*., 1994; Smith *et al*., 2004; Govindarajulu *et al*., 2005; Fellbaum *et al*., 2014). However, the effect of the nature and amount of nutrients in the compartment restricted to AM fungi, and particularly of combination of N and P, on MGR, remains to be determined.

Plant aboveground biomass can be estimated in a fast and non-destructive way by image analysis. In *B. distachyon*, this biomass was found to be well correlated with the shoot “biovolume” (Poire *et al*., 2014).

Image segmentation can be performed by chaining various image processing tools in a pipeline. Creation of pipelines and/or adjustment of each tool parameters is required to set-up segmentation of images captured by camera on phenotyping platforms. There are many freely available Image processing libraries or programs, but to our knowledge none that has a user interface, can generate pipelines that can be rapidly tested on defined image subsets and apply the analysis pipelines on large amount of image data.

Here we describe an experimental system based on a two-compartment system allowing the measurement of *B. distachyon* MGR using a state-of-the-art high-throughput phenotyping facility. We show that both N and P mobilization by the AM fungi is critical for MGR in *B. distachyon* and that there is genetic variability in *B. distachyon* for MGR. We also provide a new Python-based software tool called “*IPSO Phen*”. This open-source tool allows to configure and test image processing tools embedded in pipelines that can be executed on large amount of images generated by automated plant phenotyping facilities.

## Results

### Access of AM fungi to nitrogen and phosphorus is required for positive mycorrhizal growth response in B. distachyon

Our first goal was to design an experimental system allowing measurement of MGR in *B. distachyon* grown in pots. To control the nutrient availability for the plant and the AM fungi, we used a 1:1 (v/v) mixture of calcinated-clay and sand as substrate. To mimic the fact that *in natura* AM fungi can access nutrients that are not available to the plant roots, we use a two-compartment system. A plastic tube covered by a nylon mesh (hyphal compartment, HC), fine enough to exclude plant roots but not the AM fungal hyphae, was filled with calcinated-clay supplemented with N and/or P and placed at the bottom of 3L pots (Fig. 1a). Since N is a diffusible nutrient, part of the N can diffuse out of the HC and be accessible to plant roots. In order to limit this effect, we made sure that the total quantity of N introduced in the HC corresponds to the reported need for *B. distachyon* optimal growth (David *et al*., 2019). As P is not diffusible and would thus not leach out the HC, it was put in excess. When no N and P were supplemented in the HC, they were provided to the plants as phosphate directly mixed in the soil or nitrate through a nutritive solution.

**Figure 1.**
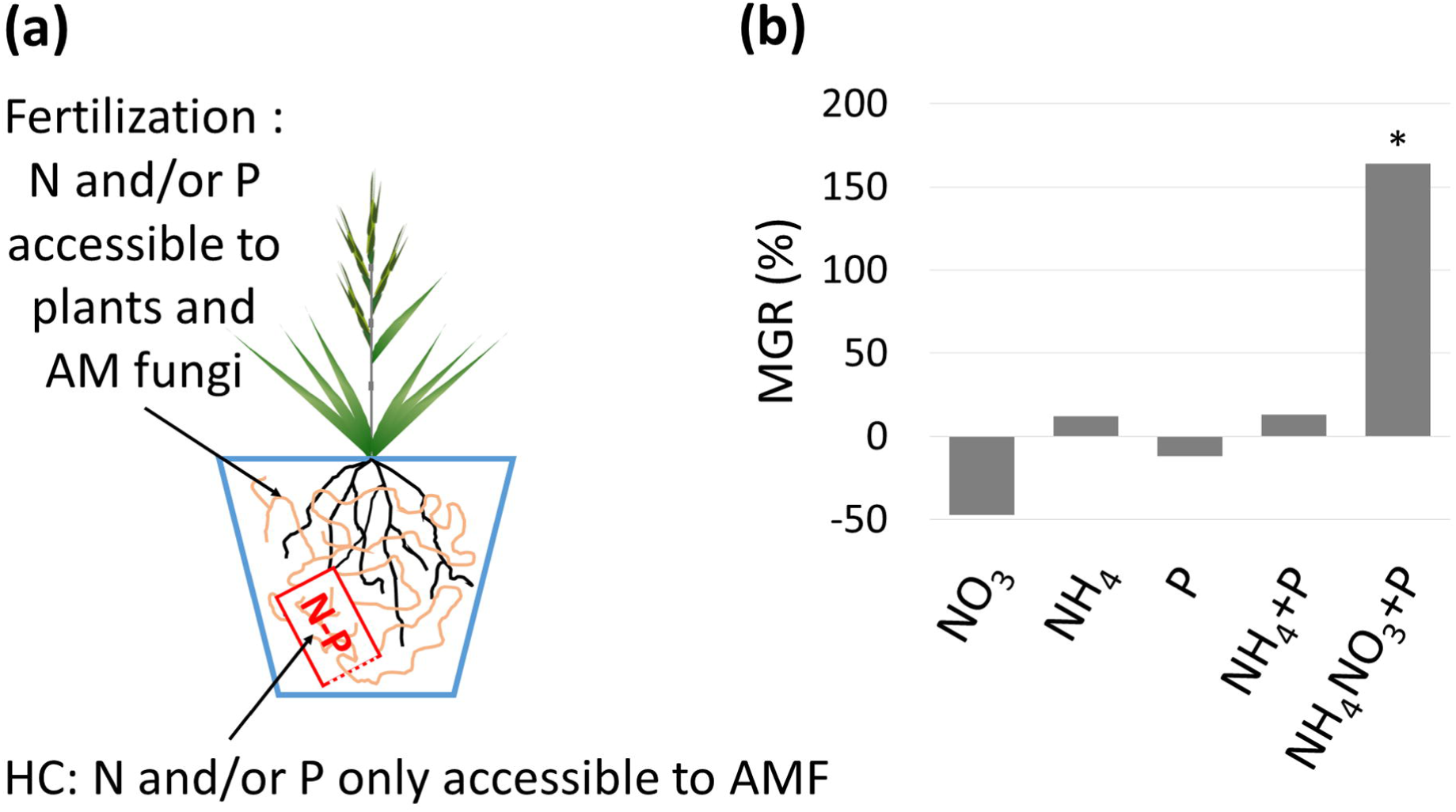
AM fungi access to N and P is required to stimulate Bd3-1 growth of in controlled conditions. a) Schematic representation of the experimental system. One plant was grown per 3L pot. A tube separated by a nylon mesh (hyphal compartment, HC, red) and containing various sources of N and/or P nutrients was placed at the bottom de the 3L pot. AM fungi (orange) but not the roots (black) accessed to the N and P in the HC. b) Indicated N or P (KH_2_PO_4_) sources were put in the HC. Plant aboveground parts were harvested 12 wpi and dry weight (DW) were measured. Mycorrhizal Growth Response (MGR) was calculated as (mean DW of inoculated plants – mean DW of non-inoculated plants) / mean DW of non-inoculated plants. * indicates a significant difference in DW between inoculated and non-inoculated plants. n=4 individuals (from one biological replicate).

In order to test the effect of N and/or P in the HC on *B. distachyon* growth, seedlings of the *B. distachyon* genotype Bd3-1 were inoculated with or without spores of the AM fungal species *Rhizophagus irregularis* and grown for 4 weeks in a growth chamber. Some plants were harvested to determine by microscopy the level of fungal colonization in roots. At 4 weeks post inoculation (wpi), in average, 32% of each root system was colonized by AM fungi. No colonization was found in non-inoculated plants. The left over plants were grown for an extra 8 weeks in an automated greenhouse (Fig. S1a). At the end of the experiment (12 wpi), in average, 56% of each root system was colonized by AM fungi. Aboveground dry weights were also measured. A significant increase in Bd3-1 aboveground biomass was observed in mycorrhizal plants only when the HC was supplemented with a mixture of phosphate, ammonium and nitrate (Fig. 1b). Similar positive MGR on Bd3-1 was observed using sand alone as a substrate (Fig. S2).

This result shows that it is possible to establish a mutualistic symbiosis between *B. distachyon* genotype Bd3-1 and *R. irregularis* in controlled conditions, but only when AM fungi have access to a source of N and P.

### “ISPO phen”, a Python-based software tool, enables a fast setup and test of image processing pipelines for optimized segmentation and feature extraction

Instead of measuring aboveground dry weight, we aimed to estimate plant aboveground biomass through image analysis. This required segmentation of the plant pixels over the background. This can be easily done for a limited number of plants using freely-available analysis tools popular among the plant phenotyping community, namely ImageJ (Rueden *et al*., 2017) and the Python based Scikit-image (van der Walt *et al*., 2014), OpenCV (Bradski. 2000) or PlantCV (Gehan *et al*., 2017). However, users are required to adjust image processing tool settings individually and tests must be done manually, which is time consuming. The procedure of setting and testing pipeline items in sets containing large amount of images can easily become cumbersome. Plant high-throughput phenotyping generates large amount of image data. Here each plant is imaged daily (8 images with a 45° angle between each imaged, Fig. S1b) over a two-month period (60 days). Considering the 264 pots as the maximal set up possible in the phenotyping greenhouse 2 of our facility (Fig. S1a), one such experiments could generate up to 8×60×264= 126 720 images. Our goal was to optimize the high-throughput segmentation of the plant pixels. Important features that we thought to be essential for this are the possibility to : (i) rapidly and semi-automatically test the solution space of given segmentation parameters, (ii) generate and test analysis pipelines, (iii) apply analysis pipelines on large amount of image data. As these features are not well served by the existing tools, we set out to develop *“IPSO phen”* a software built with Python using OpenCV, numpy (van der Walt *et al*., 2011), pandas (McKinney, 2010) and Scikit-image. *IPSO phen* also includes a user interface, access to image database and the possibility to create or import new tools, including some tools already developed in PlantCV for instance. The *IPSO phen* documentation (https://ipso-phen.readthedocs.io/en/latest/installation.html) and the source code (https://github.com/tpmp-inra/ipso_phen) are freely available.

Fig. 2a displays the flow chart that was defined and applied on the *B. distachyon* side images generated in this work. *IPSO Phen* can the save pipelines as Python scripts (see supplementary files: pipeline_script.py or binary files: pipeline_binary.tipp) that can be used to restore the process or be modified to better suite one’s needs.

**Figure 2:**
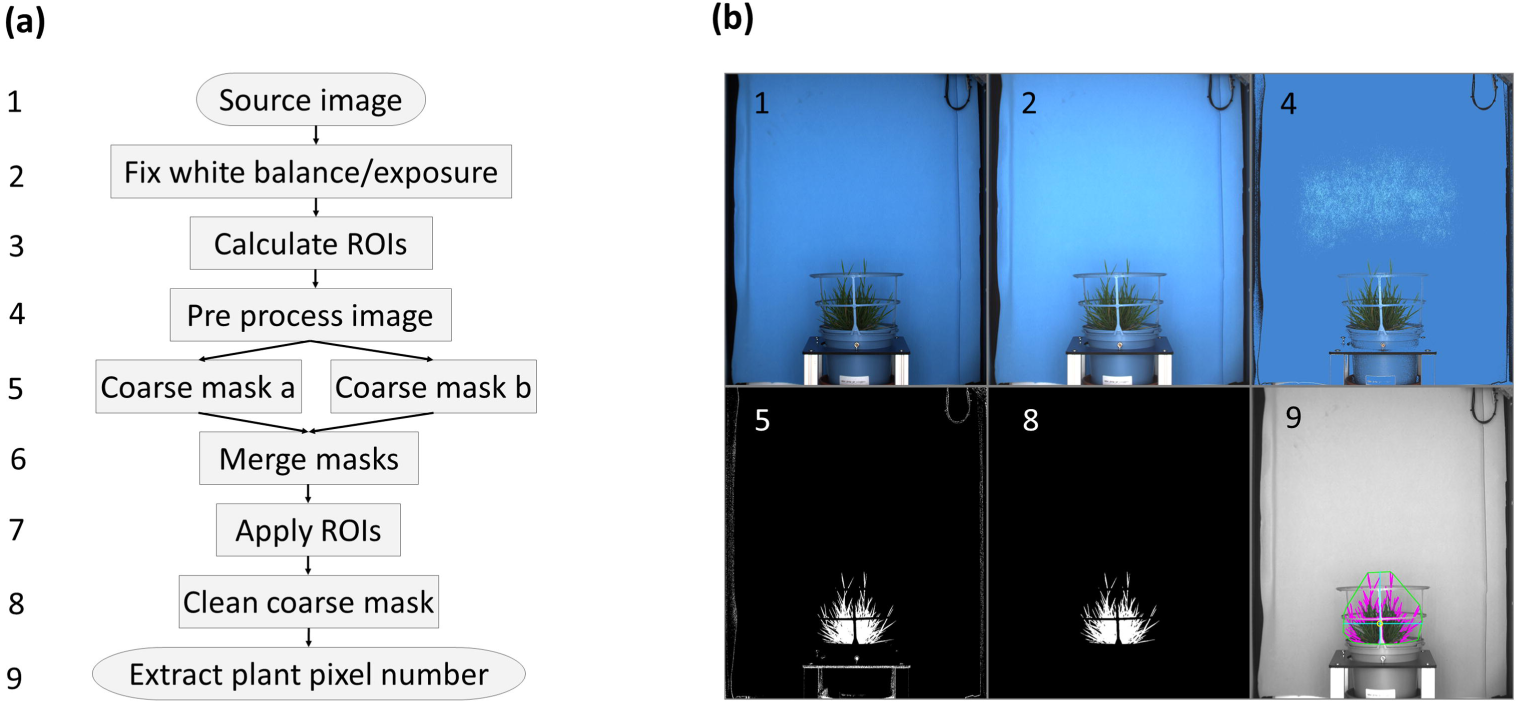
*IPSO phen* allows fast creation of pipelines to estimate *B. distachyon* aboveground biomass from side view images. a) Schematic representation of the pipeline used for *B. distachyon* image segmentation. b) Example of image processing at various steps of the pipeline (only steps 1, 2, 4, 5, 8 and 9 are illustrated). Numbers correspond to the step indicated in a). 1: **source image**, presence of a surrounding tutor cage, presence of reflection and diffraction noise. 2: **white balance and exposure.** This will fix any white balance or exposure errors that occurred during acquisition and will simplify comparisons as all images will have the same settings. 3: Regions of interest (ROI), this can be done manually or automatically, the types of ROI used here were the following: (i) keep ROI, everything outside will be deleted. (ii) safe ROI, contours inside this section won’t be treated as strictly as those outside. (iii) open ROI, a morphology open operation (erode + dilate) will be applied inside. (iv) enforcer ROI, checks that the object is where it’s supposed to be. 4: any **transformations** that renders segmentation easier, for this analysis we choose to replace the colour of darker and lighter as well as dominantly blue zones. Modification made at this step are not considered when images features are extracted 5: building **coarse masks;** in this pipeline a maximum of three coarse masks are needed: a mask to remove the tutor cage, a mask to remove the background, and another to remove the sensor noise. 6: **Merge Masks**, coarse masks as merged using logical operators “AND”, or “OR” depending on the mask. 7: **Apply ROIs**, “keep”, “delete” and “morphology” ROIs are applied at this step. 8: **Clean Coarse** mask, clean the coarse mask defined at step 5. 9: **extract features** according to the phenotyping question, here the projected plant pixels of each angle/image. For more information, see readthedocs (https://ipso-phen.readthedocs.io/en/latest/installation.html).

### Versatility of the “IPSO phen” segmentation pipeline for B. distachyon analysis

An important feature of an image analysis process is that it has to be capable of both handling large amount of data and be sufficiently versatile to cope with variation in the image features and quality. To illustrate this we choose to present the segmentation performance of the developed *IPSO phen* pipeline on small plantlets (Fig. S3a), larger and flowering plants (Fig. S3b), images with line errors (Fig. S3c) and images where misplacement of the covering foam disc (Fig S3d). The line errors are the result of data transfer issues between the camera and the database, can affect up to 10% of the images on some given days. The foam discs misplacement can be from the initial installation or appear during the experimentation and are quite rare event. Although not so common this could not be ignored and the coarse mask addressed the issue. The biomass of both small plantlets (initial stages of growth) and cases of data transfer error and disc misplacements could thus be managed by the analysis pipeline and return usable data points. The images of larger, flowering plants was not used in this work, but the pipeline could readily exploit characteristic features of this development stage.

The addition of the tutor cages on top of the pots was required to avoid any *B. distachyon* tillers from falling (which some genotype have a tendency for) but added an extra segmentation challenge, not only due to the division of plant part in sectors defined by the cage structure, but also bringing some extra light reflexion. These cages also mask plant material, with a more pronounced effect on smaller plants. In order to have the best segmentation, we applied different tools to several region of interest (ROI) of the image (step 3 of pipeline Fig. 2a).

### B. distachyon side image-determined “projected surface median” correlates with aboveground shoot biomass

We decided to calculate a proxy of plant shoot biomass by using side images. This proxy was generated for each plant imaging passage by calculating the median plant pixel number of 8 images taken at consecutive 45° angles. To determine whether this plant “median projected surface” (MPS hereafter) provided by image analysis correlate with the plant biomass over the *B. distachyon* growth cycle, we harvested *B. distachyon* plants weekly and measured both the MPS from images and the actual biomass. We found a good correlation (R^2^=0.975) between the aboveground MPS and the aboveground dry weight (Fig. 3). The correlation was lost once the spikelets appeared (Fig. S4), showing that the MPS is a good approximation of shoot but not seed biomasses.

**Figure 3.**
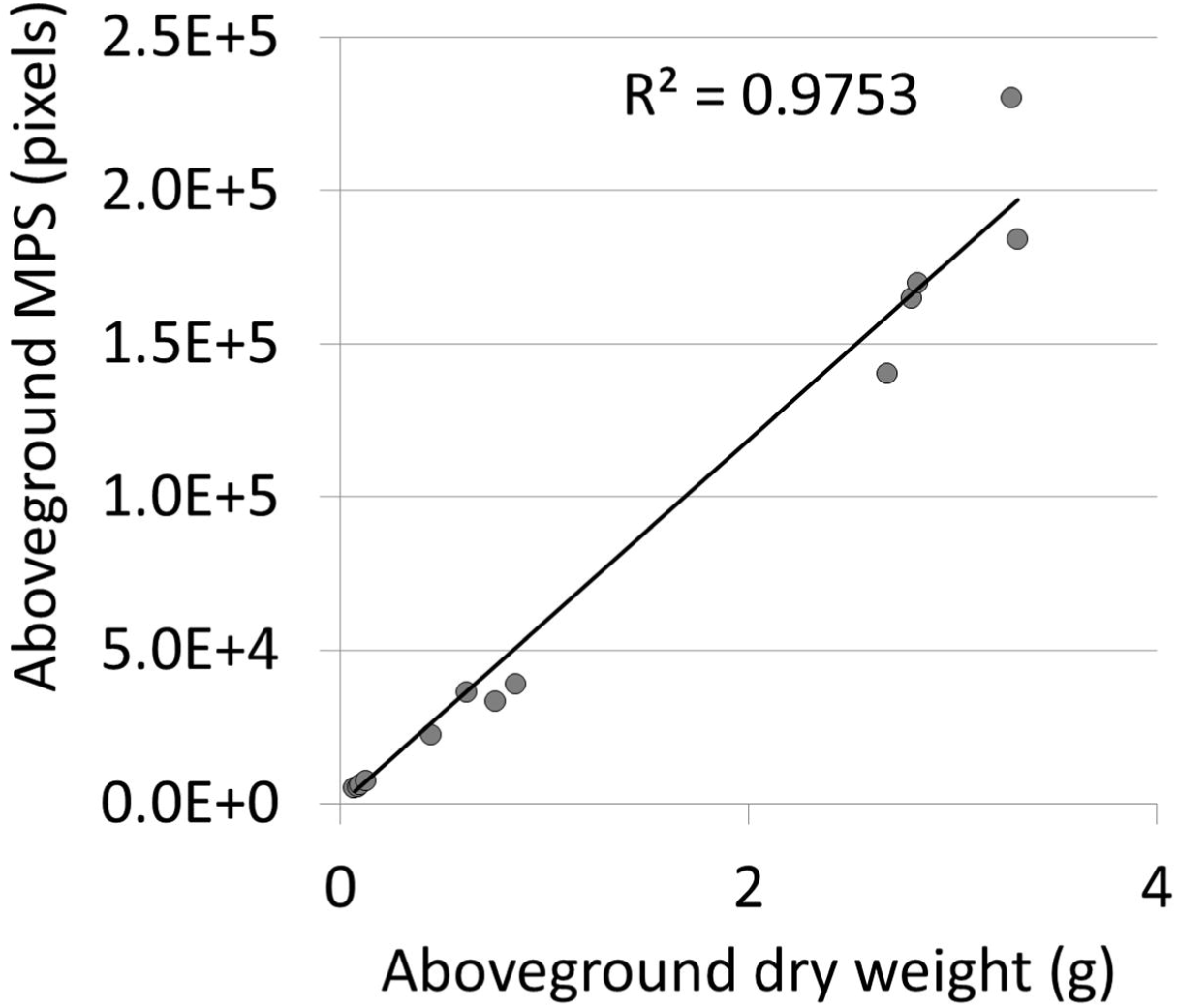
*B. distachyon* aboveground shoot biomass can be estimated by side image segmentation. Plot of the aboveground MPS and dry weights. The MPS is the median of the plant pixel number obtained from the 8 images taken with 45° angles. Plants were harvested weekly and the MPS was determined on images taken immediately before harvesting.

### Image analysis shows that mycorrhizal growth response in B. distachyon depends on plant nutrition through AM fungi

We then applied image segmentation and MPS calculation to follow the growth kinetic of Bd3-1 in presence or absence of AM fungi (Fig. 4a). To quantify the effect of AM fungi on growth, we measured the “area under the curve” (AUC) for each individual (Fig. 4b) we also calculated the MGR using the MPS at the end of the cycle (Fig. 4c). Better plant growth was observed in presence of AM fungi when N and P were added in the HC and plant N/P fertilization was low. In this condition, a significant difference in the AUC was found between mycorrhizal and the non-mycorrhizal plants (Fig. 4b), leading to a positive MGR (Fig. 4c). Analysis of plant MPS at each day of the kinetic showed that a significant difference between mycorrhizal and the non-mycorrhizal plants appeared 45 days after inoculation with AM fungi (17 days after addition of the HC in the experimental system). In contrast, no significant increase in growth of mycorrhizal plant compared to non-mycorrhizal plants, was observed when no N and P were added to the HC and N/P fertilization was low (Fig. 4a-c). Similarly, when N and P were added to the HC but plants were fertilized with high levels of N and P, no effect of AM fungi on plant growth was observed (Fig. 4a-c).

**Figure 4.**
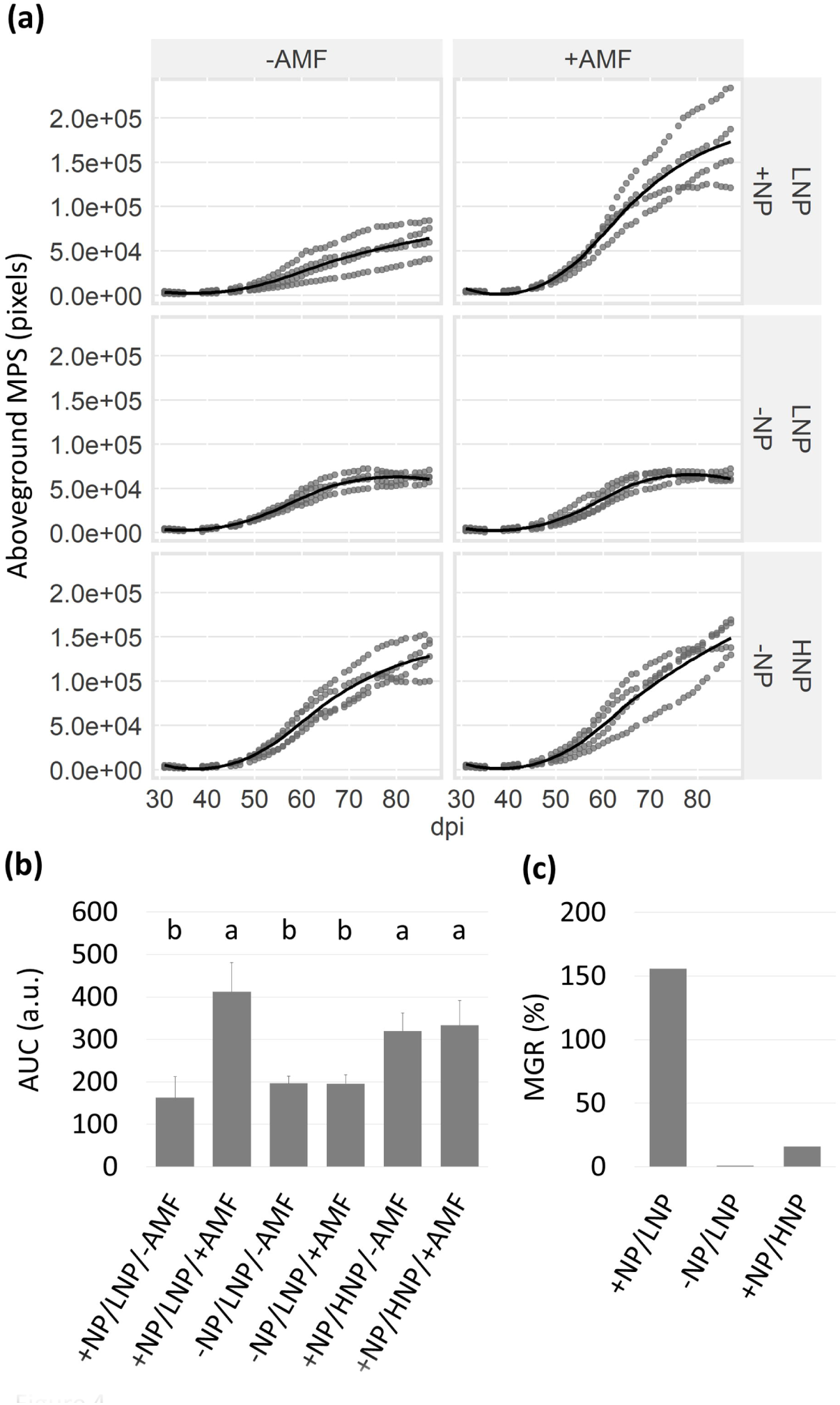
Image segmentation allows to measure mycorrhizal growth response in Bd3-1. a) Growth curves of Bd3-1 in presence (+AMF) or absence (-AMF) of *R. irregularis* grown in pots with HC containing N and P (+NP) or not (-NP). Plants were fertilized with low levels of N and P (LNP) or high levels of N and P (HNP). n=4 individuals (from one biological replicate). b) Differences in growth were quantified as areas under the curves (AUC). Different letters represent statistically different categories. c) Mycorrizal Growth Response (MGR) was calculated using the plant MPS at the end of the growth as (mean MPS of inoculated plants – mean MPS of non-inoculated plants) / mean MPS of non-inoculated plants.

To determine whether there is genetic variability in *B. distachyon* for MGR and whether our experimental conditions allows to measure it, we initiated a screen *B. distachyon* genetic diversity for MGR. Differences were observed between the 16 genotypes tested ranging from positive to negative MGR (Fig. 5).

**Figure 5.**
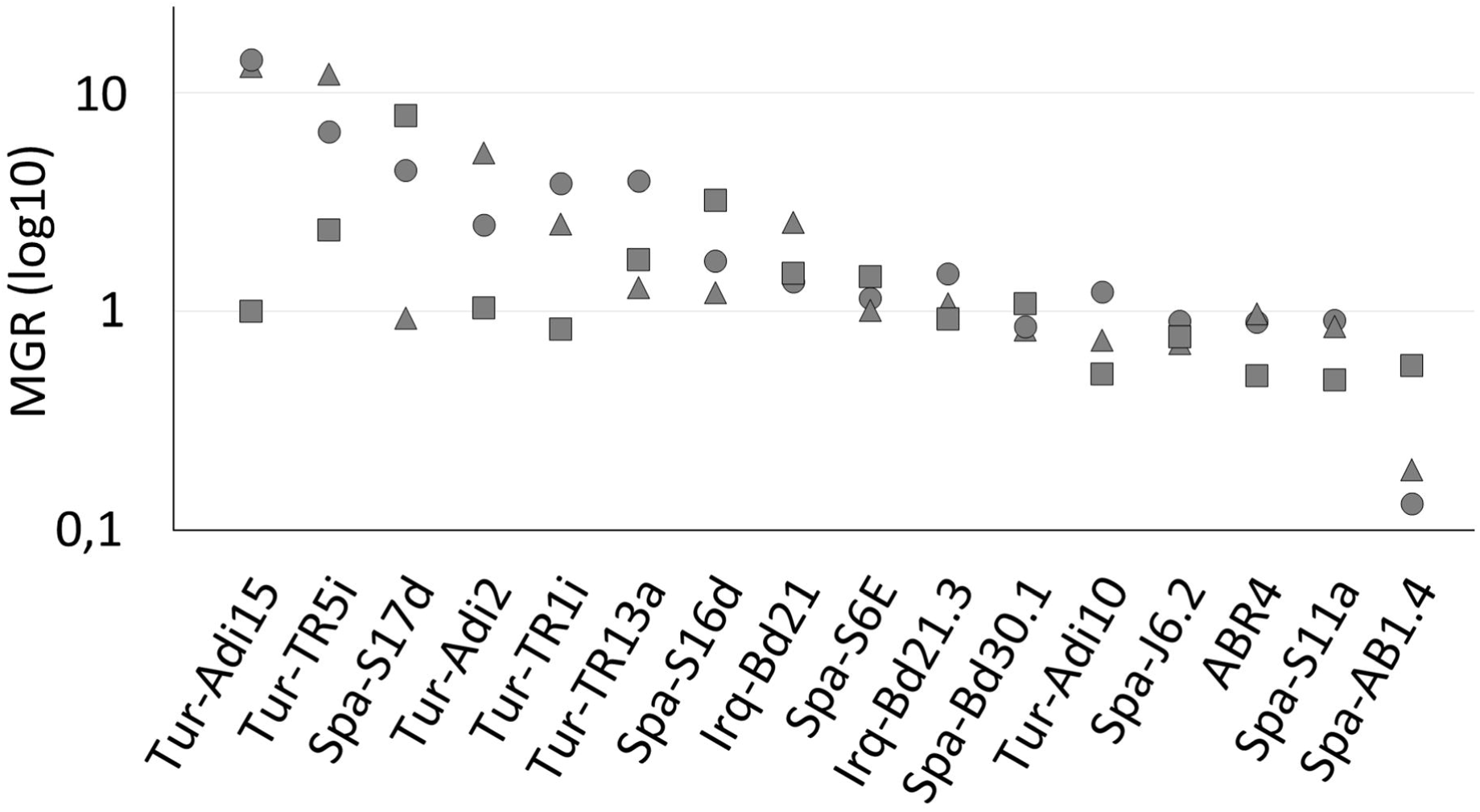
Image segmentation allows to screen *B. distachyon* genetic variability for mycorrhizal growth responses. Plants were grown in presence or absence of *R. irregularis* grown in pots with HC containing N and P and were fertilized with low levels of N and P. n=3 individuals (from 3 biological replicates). Mycorrizal Growth Response (MGR) was calculated as log10 (MPS of inoculated plants / MPS of non-inoculated plants) in each biological replicate using the plant MPS at the end of the growth. Triangles, Dots and squares represent MGR in biological replicate 1, 2 and 3 respectively.

## Discussion

Plant automated high throughput phenotyping enables to have more data points for a plant trait of interest compared to classical human-recording phenotyping. These can be more individuals as well as more time points. Moreover, the automated phenotyping platform, including the watering facility, allows to decrease the experimental variability within a given experiment. We would like to emphasis that we observed in this work a very low growth variability between plants with the same treatment within each replicate. Although not quantifiable as such, this variability was much lower compared to what we observed in various experiments we ran previously in growth chambers or greenhouses (although the plants were not grown under the exact same condition). However, we observed a strong variability between the first/second and the third replicates when we analyzed the *B. distachyon* genetic variability for MGR. For an unexplained reason, in the third replicate all genotypes started to flower during the 4 weeks the plants were in the growth chamber while the first spikelets started to appear 1 week after transfer in the greenhouse in the two first replicates. This had a strong effect on MGR although the type of effect (positive or negative MGR) was similar between the replicates. Moreover, the early flowering plants had a reduced growth that affected the quality of image-segmentation with the imaging parameters we selected for our experiments (Fig. S6). Low variability of growth between individuals was also observed previously and led to the successful phenotyping of other plant-microbe interactions (Mazo-Molina *et al*., 2019). We thus believe that the plant rearing conditions in the TPMP facility (Fig. S1a) are quite homogeneous, minimizing the biological variability.

We chose to estimate the shoot biomass only using side images, and not a combination of side and top images like (Poire *et al*., 2014). This was mainly due to a phenotyping time consideration, as the top and side imaging can’t run simultaneously (lighting conditions are different). Top imaging would have implied a second phenotyping job, not compatible with other experiments running on the facility at the same time. However, the correlation with biomass was similar to that found in (Poire *et al*., 2014). Moreover, the use of the median pixel plant number has the advantage of being quite tolerant to accidental variations (for instance a tiller that would be blocked in the tutor cage).

### IPSO Phen a new opensource software allowing the fast development of pipelines for plant image segmentation

In this work we have developed a new software suite that we believe is well suited to the analysis of large amount of image data, typically generated by high throughput plant phenotyping. The most popular open-source solutions for analyzing large plant-phenotyping data is PlantCV (Gehan *et al*., 2017). The clear advantages of PlantCV are (i) the fact that it is built on known image-analysis toolkits like OpenCV or Scikit-image, whilst being (ii) easier to use than OpenCV (iii) its opensource nature, and (iv) large user community. We believe that *IPSO phen* could be a good addition/companion to PlantCV. Indeed *IPSO phen* fills the gap of known drawback of PlantCV as it provides (i) a User Interface, (ii) doesn’t require one to be a Python programmer, (iii) provides the possibility to build and test image processing pipelines as they are being built. Like PlantCV, *IPSO phen* is built on industry standards (OpenCV, Scikit-image, numpy and pandas), it is easily extendable as new tools and plugins can be created and/or added from known sources. We have imported popular PlantCV tools within the tool kit available in *IPSO phen*, these tools (by convention; named PCV in *IPSO phen*), can readily be integrated and tested in any new pipeline. A grid search, allows to explore a wide solution space in a very simple way, with easy inspection and selection by the user of the desired tool settings. Finally, in our experiments, imaging was performed daily and the raw images where stored in a dedicated database (not discussed in this work). *IPSO phen* software also has the advantage to directly access the desired images in the database. In conclusion, *IPSO phen* is meant to be used for high throughput as it connects directly with image databases (or filesystem), the pipeline functions can be tested on fixed or random sets of images at every step and the mass processing can be done in various steps.

*IPSO phen* is fully functional, but still in its early development. A roadmap for the future would be an improvement of the User Interface and a more flexible and intuitive pipeline builder.

### Co-transport of N and P might be required for efficient mycorrhizal growth response in B. distachyon

In our controlled conditions, *B. distachyon* growth stimulation by AM fungi was observed only when N and P were added in the HC accessible to the AM fungi. This suggests that the growth stimulation is due to plant N and P nutrition through the AM fungi. It has to be noted that the growth of *B. distachyon* was similar when N and P were transported through AM fungi (+ AM fungi, fertilization with low N and P in the pot) or directly acquired by the root system (-AM fungi, fertilization with high N and P in the pot). The lack of *B. distachyon* growth increase between the conditions in which N and P were added or not in the HC in absence of AM fungi suggests that diffusion of at least one the two nutrients is limited. The requirement of the presence of both N and P in the HC for positive MGR in *B. distachyon* was a surprising result. Indeed, positive MGR on maize was observed by introducing only P to the HC (Gerlach *et al*., 2015). However, it might explain the absence or negative MGR observed in many experimental systems used in controlled conditions even if P transport from the HC to the plants was measured (Smith *et al*., 2004; Li *et al*., 2006). In our experimental system, when either only N or P were added in the HC, the other nutrient was added to the substrate outside of the HC and thus accessible to both AM fungi and roots. The lower plant growth compared to the condition in which both nutrients were added to the HC or directly in the substrate, suggests that uptake both nutrients is inefficient if not performed through the same mechanism (direct uptake or through AM fungi). It has been hypothesized that N and P can be co-transported in the AM fungal mycelium from the place they are collected to the arbuscules in plant roots. N is transported as arginine and P as polyphosphate, and arginine can bind to polyphosphate (Govindarajulu *et al*., 2005). Stoichiometry of N and P in the mycelium might thus affect efficiency of their transport. Alternatively, colonization of plants by AM fungi can strongly repress expression of phosphate, nitrate and ammonium transporters involved in direct nutrient uptake, including in *B. distachyon* (Hong *et al*., 2012). The P availability in the soil also interferes with N uptake and AM-dependent regulation of N transporters (Nouri *et al*., 2014). It could be that colonization by AM fungi induces repression of direct nutrient uptake even if the AM fungi cannot provide all limiting nutrients for plant growth. It should be determined to which extent availability of N, P or N+P in the HC affects the direct uptake of both N and P.

Genes coding for both nitrate and ammonium transporters are found in AM fungi (reviewed in Chen *et al*., 2018). When in competition with nitrate, ammonium (provided as ammonium nitrate in the HC) appeared to be the preferred N source for AM fungi, both in terms of uptake by AM fungi and delivery to plants (Tanaka & Yano, 2005). However, when only nitrate or ammonium were used as N sources (both for plant and AM fungi), better N transport by AM fungi to plants was obtained with nitrate (Hawkins & George, 2001). The authors suggested that pure ammonium can be deleterious for AM fungi. In our experimental conditions, we found a positive MGR if ammonium and nitrate were added together with P in the HC but not if only ammonium was added together with P, suggesting that a balance between ammonium and nitrate is required for efficient N uptake by AM fungi.

We observed positive MGR on the *B. distachyon* genotype Bd3-1 from 45 days after inoculation and 17 days after addition of the HC. Once the HC was added to the pots, it might take a few days for AM mycelium to reach the HC and start to uptake the nutrients it contains. We have not observed significant growth depression in the mycorrhizal plants before the AM fungi reach the HC and can provide a nutritional benefit. This suggests that in our conditions the carbon cost of AM establishment is not limiting for plant growth or is compensated by an increase of plant photosynthetic activity.

Here we show that AM fungi can stimulate the increase of *B. distachyon* shoot biomass. It would be interesting to determine which other plant traits are affected during AM. Image analysis can easily determine additional aboveground plant traits such as height, width, and various features of the aboveground plant shape. Automatic analysis of other traits such as the tiller number or the flowering time would require development of new pipelines/tools. In addition, seed development would be another important trait to follow. It would be interesting to analyze whether any changes in imaging variables correlate with effects of AM fungi on seed quantity and/or quality and could be used as proxy to follow effects of AM on seed production.

Testing MGR in 16 *B. distachyon* genotype, we observed variability between genotypes, ranging from a positive to negative MGR. Reasons that can explain host genetic variability for MGR are not known. Differences in host efficiency to acquire N and P from AM fungi could be one of them. Alternatively, AM fungi induce plant defense (Güimil *et al*., 2005; Campos-Soriano *et al*., 2012; Watts-Williams *et al*., 2019) and trade-off between plant growth and defense are known (reviewed in Karasov *et al*., 2017). Differences in level of host defense induction by AM fungi might explain difference in MGR. The genetic variability observed in *B. distachyon* is similar to that observed in other species except in maize for which only positive MGR were found (Kaeppler *et al*., 2000; Sawers *et al*., 2017). It is interesting to note that in maize, positive MGR was also found when only P was added in the HC. Together it suggests that both nutrition (direct nutrient uptake and/or through AM fungi) and MGR are different in maize and *B. distachyon* or other grass species.

Identification of plant loci/genes controlling MGR would be of interest to help breeding for optimized MGR. The experimental design we setup, the genetic variability we found in *B. distachyon* and the software we developed for high throughput image analysis make feasible analysis of MGR on large panels of *B. distachyon* accessions / recombinant lines and quantitative genetics allowing identification such gene/loci in this species.

## Materials and methods

### Plant growth conditions

*Brachypodium distachyon* Bd3-1 was used for setting up the experimental system. Variability of mycorrhizal growth responses was tested on the other indicated genotypes. Seeds were surface sterilized for 30 sec in 70% ethanol and for 5 minutes in 3.2% active chlorine bleach solution. Seeds were placed in agar plates and incubated at 1 week (Bd3-1, Bd21, and Bd21-3) or 4 weeks (the other genotypes) at 4° C. The seeds were germinated for 3 days at 25°C before planting.

Seedlings were planted in peat pots (Fig. S7a) filled with about 250 ml of a 1:1 (v/v) mix of calcinated clay (Attapulgite Sorbix) and quartz sand (0.7-1.3 mm) and inoculated with 2000 spores of *Rhizophagus irregularis* DAOM 197198 (Agronutrition, France).

Plantlets were grown for 4 weeks in a growth chamber (16h of light, LED light at 300 μmol.m^-2^.s^--1^, 24 °C - 8h of dark, 18 C). Each pot was individually watered with the same volume of a low N and P nutritive solution 3 times per week (Table S1). After 4 weeks, each peat pot was placed in a 3L plastic pot (3LSX, Bleu EK, Soparco, France) filled with the same substrate. A 40 ml plastic container filled with the calcinated clay and covered by a 30 μm nylon mesh was used as hyphal compartment (HC). The HC was supplemented with 305mg of NH_4_Cl, 577mg of KNO3, or 130 mg of KNO_3_+177 mg of NH_4_NO_3_ (each condition corresponding to 80 mg of N) and/or 115 mg of KH_2_PO_4_. The HC was placed at the bottom of the 3L pots. For the high N and P condition, 25ml of the N solution (10 mM KNO_3_ and 10 mM Ca(NO_3_)_2_) were added weekly and/or 115mg of KH_2_PO_4_ were mixed with the substrate used to fill the 3L pots, once the 3L pots were loaded on the automated greenhouse. A blue foam disc (eva/rubber composition, 210 g.m^-2^, 2mm thick, uniform blue colour) was glued on the rim of the pots to facilitate image segmentation (Fig. S7b). These discs also have the advantage of guiding the watering flow towards the center of the pot and lowering water loss by soil evaporation. A blue tutor cage (Sopafix 19 BAS, Bleu EK, Soparco, France) was attached on each pot to maintain all plant tillers upwards and thus allow more reproducible plant imaging (Fig. S7c). Plants were grown on the automated greenhouse (Fig. S7d) with watering three times per week with 8 ml of 2.5x nutritive solution (Table S1). Additional water was added to maintain each pot at a fixed weight corresponding to 70% of the maximal water retention capacity.

For measuring the correlation between image analysis and aboveground dry weight, Bd3-1 seedling were planted on potting soil (SB2, Proveen, The Netherlands) supplemented with 1.7g/L of osmocote (12-7-19+TE) and grown on the automated greenhouse. Four plants were harvested weekly.

### IPSO phen

The documentation files are available through readthedocs (https://ipso-phen.readthedocs.io/en/latest/installation.html) and contains all the details and explanation of how to install and use *IPSO phen*. The software was built using OpenCV (Gehan *et al*., 2017), numpy (van der Walt *et al*., 2011), pandas (McKinney, 2010) and Scikit-image (van der Walt *et al*., 2014). The source files are available through Github (https://github.com/tpmp-inra/ipso_phen). *IPSO phen* was created to (i) have a user interface, (ii) build for high throughput (image database connection, functions can be tested at each step, mass process in various steps) (iii) new tools/plugins can be added or built (declarative UI building, callbacks handled behind the scene, documentation and docstrings built automatically), (iv) Grid search to explore solution space in single process. Specific inquiries can be made using the dedicated email address

### Data analyses

Growth curve regressions were calculated using geom_smooth from R’s tidyverse library with method “loess” and “y ∼ x” formula. Areas under the curves were measured using the script audpc in the R library agricolae. Statistical differences in the aboveground dry weights, plant MPS at the fixed days or AUC were analyzed using a Kruskal Wallis test (p<0.05).

## Supporting information

SuppFigures

pipeline script

## Acknowledgements

Seeds were kindly provided by Richard Sibout (INRA, France) and John Vogel (DOE-JGI, USA).

## Author contributions

BL and NP designed and planned the experiments. FMM designed and created *IPSO phen* software. CR, LB, MG, MK, FD performed the experiments. FMM, NP and BL analyzed the data. FMM, NP and BL wrote the manuscript.

## Funding information

This work was supported by the “Laboratoire d’Excellence (LABEX)” TULIP (ANR-10-LABX-41) and by the Institut carnot “Plant2Pro” (contracts STRESSNSYM and PHENOR).

## Supporting Information

**Fig. S1**. Phenotyping system design.

**Fig. S2.** AM fungi access to N and P is required to stimulate Bd3-1 growth of in controlled conditions.

**Fig. S3**. *IPSO phen* pipeline versatility.

**Fig. S4.** *B. distachyon* seed biomass cannot be estimated by side image segmentation.

**Fig. S5**. Growth of 16 *B. distachyon* genotypes in presence or absence of *R. irregularis*.

**Fig. S6**. Normalized growth of 16 *B. distachyon* genotypes in presence or absence of *R. irregularis*.

**Fig. S7.** Experimental system design.

**Supplementary file 1:** pipeline_script.py

**Supplementary file 2:** pipeline_binary.tipp

