## Supplementary material for "Experimental system and image analysis software for high throughput phenotyping of mycorrhizal growth response in *Brachypodium distachyon*": SuppFigures

(a)

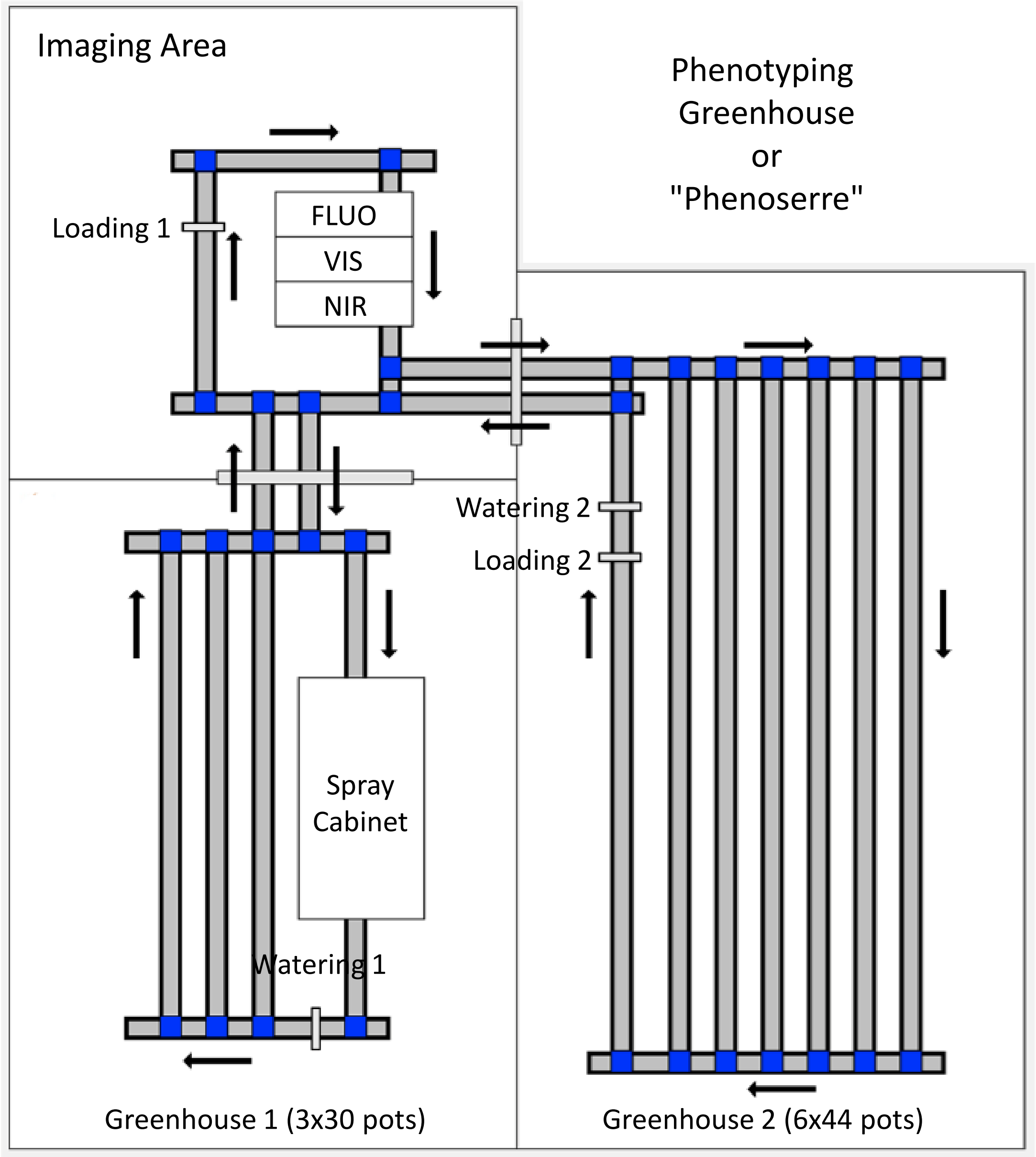


(b)

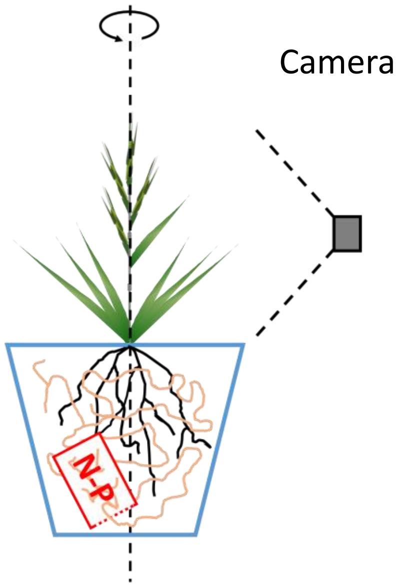


**Fig. S1. Phenotyping system design.** a) The phenotyping infrastructure to rear and image plant in the 3L pot setting. All the experiments in this work where performed in “Greenhouse 2”, planted pots were loaded (“Loading 1) into the system on individual conveyors, watered daily according to the specified protocol (Watering 2 station) and imaged daily solely using the side camera in the visible (VIS) spectrum of the “Imaging Area”. Not relevant for this work, but detailed on this scheme, are: the Greenhouse 1 and Spray Cabinet as well as the other imaging modalities: fluorescence (FLUO) and near-infrared (NIR) top and side as well as the VIS top cameras. Arrows indicate the direction of plant movement along the conveyor belts. b) Pot rotation axis and camera position during image acquisition. Each plant is imaged by rotating the conveyor at 45° 8 times.

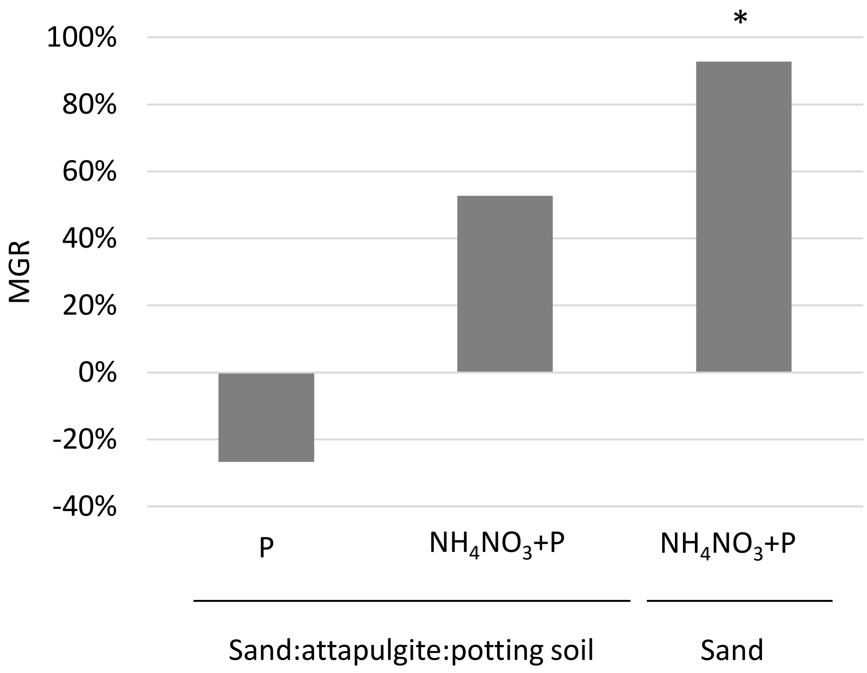


**Fig. S2. AM fungi access to N and P is required to stimulate Bd3-1 growth of in controlled conditions.** N ($NH_4NO_3$+$KNO_3$) and/or P ($KH_2PO_4$) were put in the HC. Plants were grown in a 2:2:1 mix sand:attapulgite:potting soil (motte20, Proveen, The Netherlands) or in sand. Aboveground parts were harvested 12 wpi and dry weights (DW) were measured. Mycorrhizal Growth Response (MGR) was calculated as (mean DW of inoculated plants – mean DW of non-inoculated plants) / mean DW of non-inoculated plants. * indicates a significant difference in DW between inoculated and non-inoculated plants. n=4 individuals (from one biological replicate).

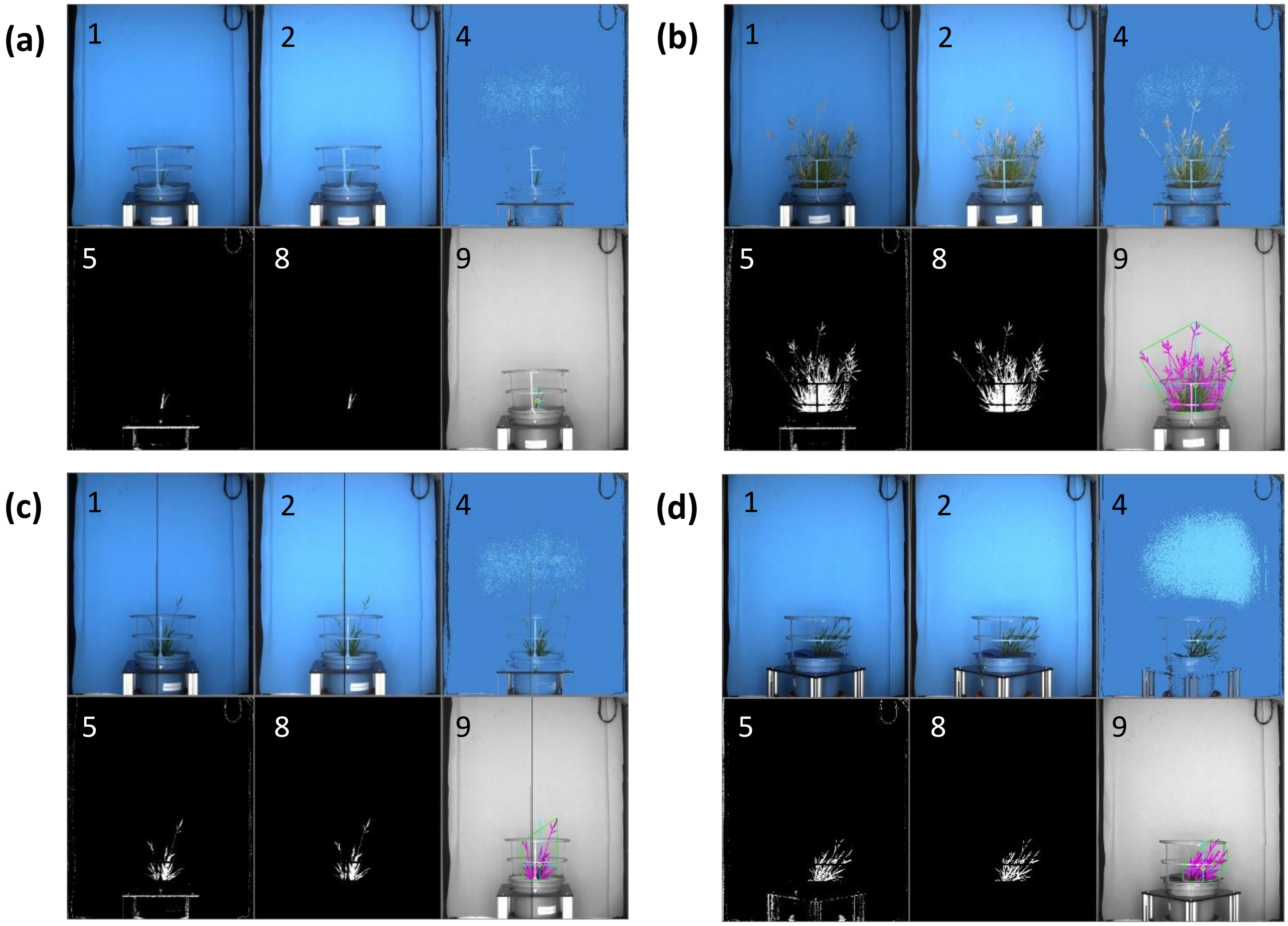


**Fig. S3. *IPSO phen* pipeline versatility.** Three examples of versatility of the *IPSO phen* developed analysis pipeline: a): small plantlets, b): large flowering/seed setting plants, c): images with line errors, d) images with foam disc misplacement. The order images in the mosaics and the legends are the same as in Fig. 2.

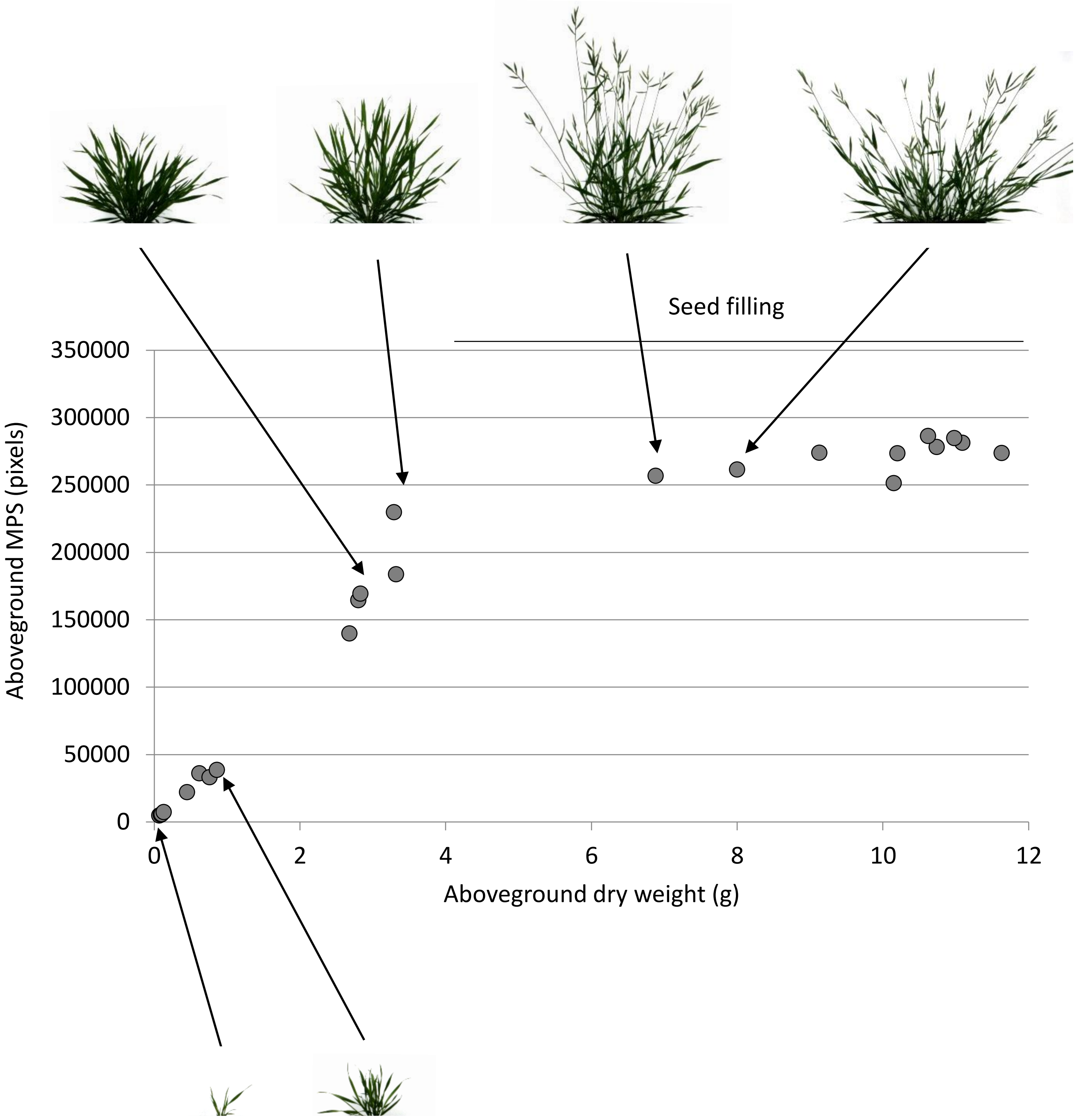


**Fig. S4. *B. distachyon* seed biomass cannot be estimated by side image segmentation.** Plot of the aboveground MPS and dry weights (same than in Fig. 3). The MPS is the median of the plant pixel number obtained from the 8 images taken with 45° angles. Plants were harvested weekly and the MPS was determined on images taken immediately before harvesting.

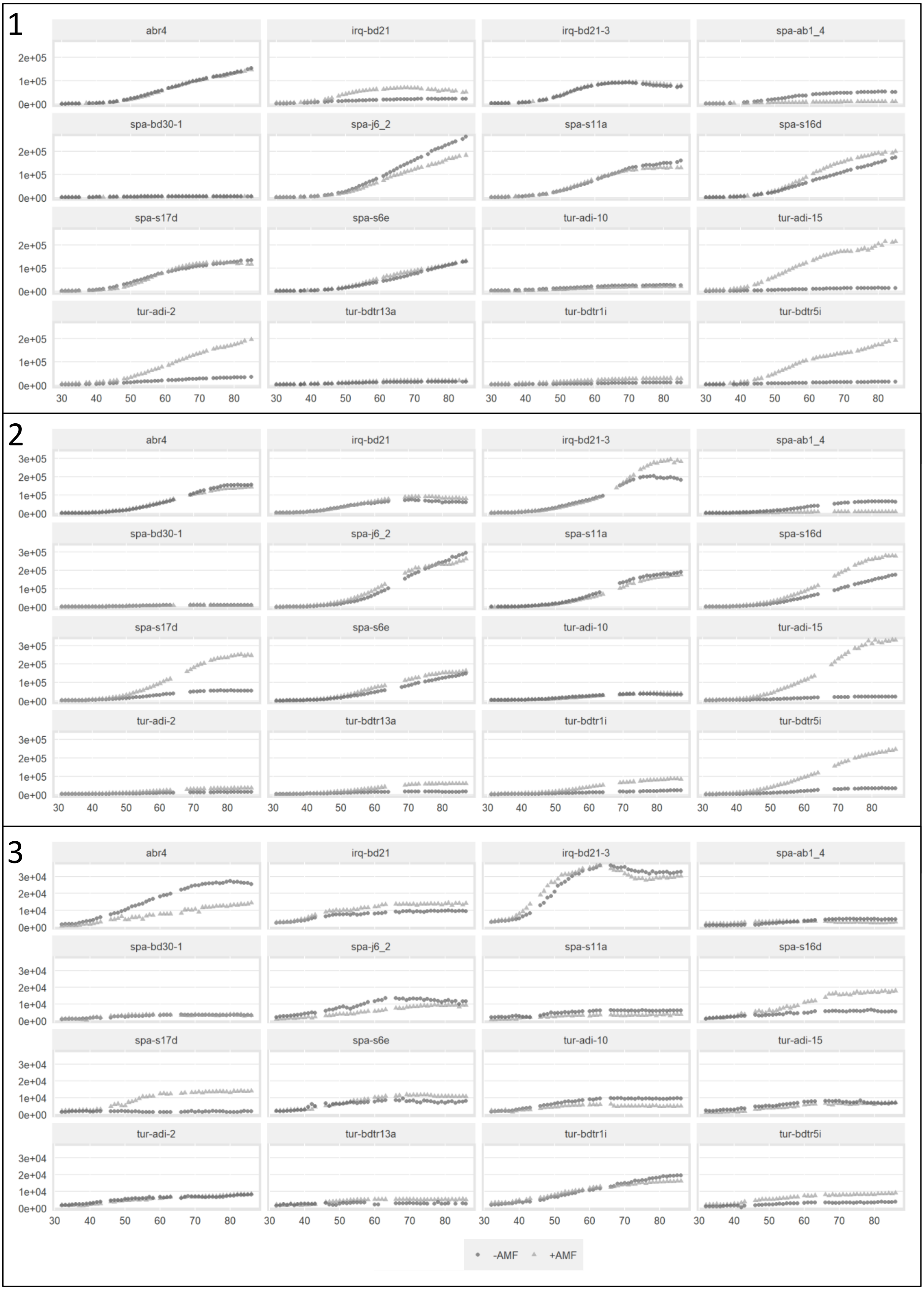


**Fig. S5. Growth of 16 *B. distachyon* genotypes in presence or absence of *R. irregularis*.** Plants were grown in pots with HC containing N and P and were fertilized with low levels of N and P. The X axis corresponds to days post inoculation with *R. irregularis* (+AMF) or without inoculation (-AMF). The 3 biological replicates are shown.

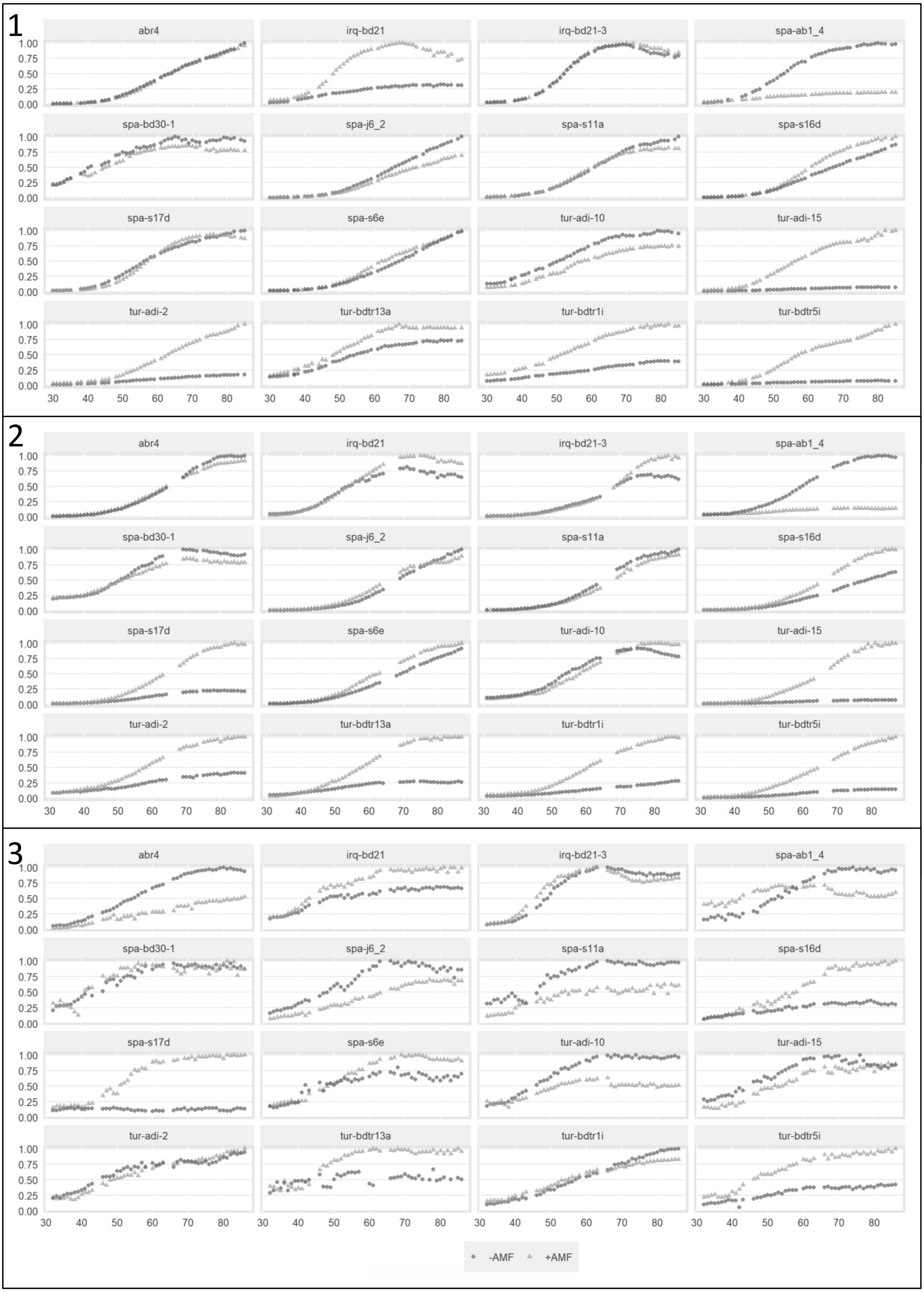


**Fig. S6. Normalized growth of 16 *B. distachyon* genotypes in presence or absence of *R. irregularis***. Same plants as in Fig. S5 except that the growth curves from were normalized by the maximum MPS measured for each genotype in each replicate (arbitrary unit, A.U.).

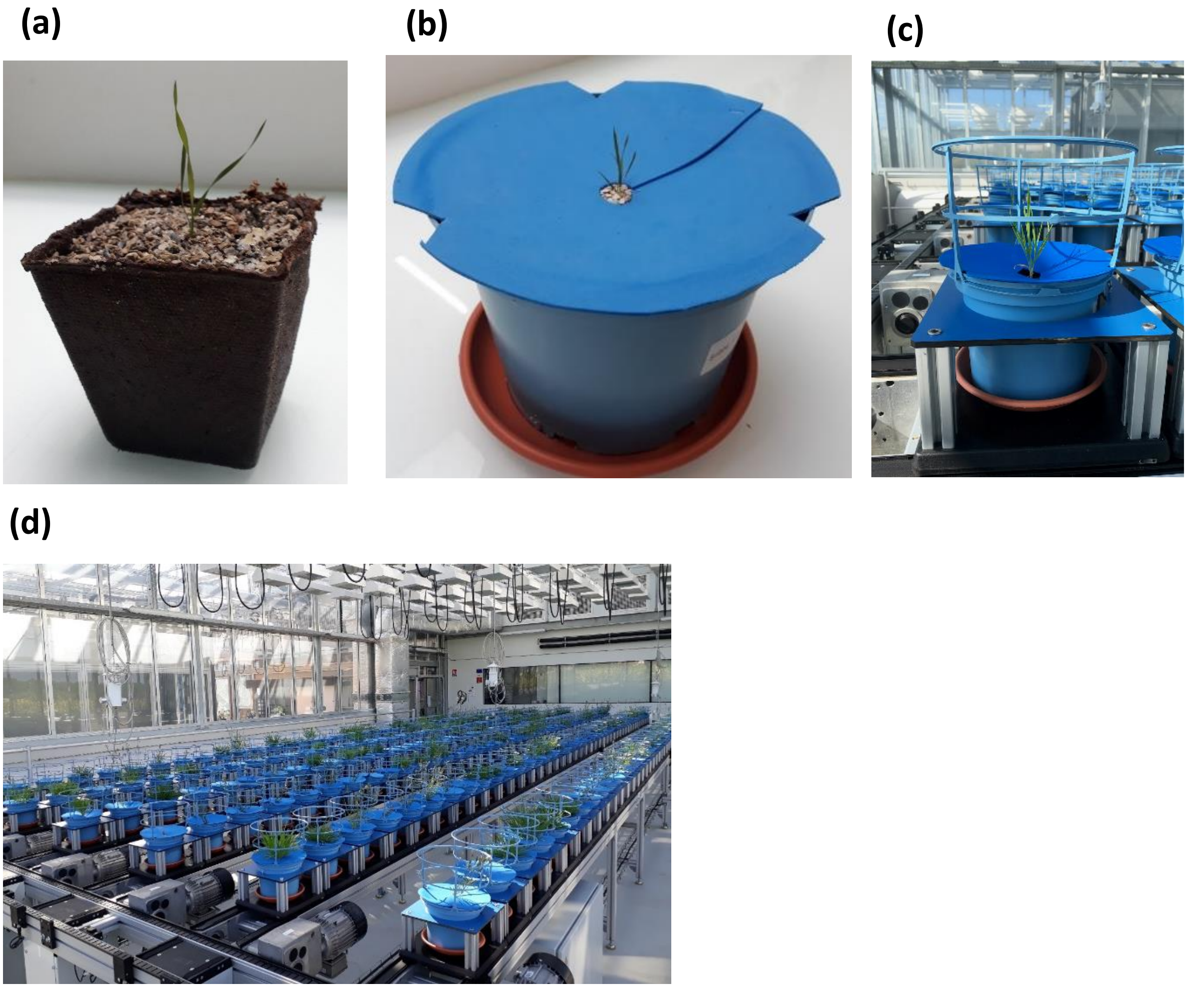


**Fig. S7. Experimental system design.** a) Peat pots filled with 1:1 (v/v) mix of calcinated clay and sand. b) 3L plastic pots filled with the same substrate. A blue foam disc was glued on the rim of the pot to facilitate image analysis. c) A blue tutor cage was attached on each pot to maintain all plant tillers in the camera frame. d) Overall view of the automated greenhouse.

| | Final concentration (µM) |
|---|---|
| Macroelements | |
| $KH_2PO_4$ | 300 |
| $Ca(NO_3)_2$, $4H_2O$ | 500 |
| $MgSO_4$, $7H_2O$ | 2000 |
| $CaCl_2$, $2H_2O$ | 3000 |
| KCl | 5000 |
| Microelements | |
| $H_3BO_3$ | 48.52 |
| NaCl | 85.56 |
| $CuSO_4$, $5H_2O$ | 1.00 |
| $MnSO_4$, $H_2O$ | 10.06 |
| $ZnSO_4$, $H_2O$ | 1.04 |
| $(NH4)_6Mo_7O_{24}$, $4H_2O$ | 0.07 |
| Na-Fe(III)-EDTA | 60 |

Table S1. Low N and P nutritive solution used to fertilize the plants
