## Supplementary material for "Experimental system and image analysis software for high throughput phenotyping of mycorrhizal growth response in *Brachypodium distachyon*": pipeline script: pipeline_script.py

```
import argparse
import csv
import cv2
import numpy as np
import os
import sys

abspath = os.path.abspath(__file__)
fld_name = os.path.dirname(abspath)
sys.path.insert(0, fld_name)
sys.path.insert(0, os.path.dirname(fld_name))

from ip_base.ip_abstract import AbstractImageProcessor
from ip_base.ipt_functional import call_ipt, call_ipt_func
from tools.csv_writer import AbstractCsvWriter


def main():
    # Get the file
    # ____________
    # Set working folder
    old_wd = os.getcwd()
    abspath = os.path.abspath(__file__)
    fld_name = os.path.dirname(abspath)
    os.chdir(fld_name)

    # Construct the argument parser and parse the arguments
    ap = argparse.ArgumentParser()
    ap.add_argument("-i", "--image", required=True, help="Path to the image")
    ap.add_argument("-d", "--destination", required=False, help="Destination folder")
    ap.add_argument("-p", "--print_images", required=False, help="Print images, y or n")
    ap.add_argument("-m", "--print_mosaic", required=False, help="Print mosaic, y or n")
    args = vars(ap.parse_args())
    file_name = args["image"]
    print_images = args.get("print_images", "n") == "y"
    print_mosaic = args.get("print_mosaic", "n") == "y"
    dst_folder = args.get("destination", "")

    # Restore working folder
    os.chdir(old_wd)

    # Build wrapper
    # _____________
    wrapper = AbstractImageProcessor(file_name)
    wrapper.lock = True
    wrapper.store_image(wrapper.current_image, 'true_source_image')
    if print_images or print_mosaic:
        wrapper.store_images = True
    if print_images:
        wrapper.write_images = 'plot'
    if print_mosaic:
        wrapper.write_mosaic = 'plot'

    # Fix exposure
    # ____________________
    wrapper.current_image = call_ipt(
        ipt_id="IptLinearTransformation",
        source=wrapper,
        return_type="result",
        method='alpha_beta_target',
        target_brightness=150
    )

    # Store image name for analysis
    wrapper.store_image(wrapper.current_image, "exposure_fixed")
    analysis_image = "exposure_fixed"

    if print_mosaic:
        wrapper.store_image(wrapper.current_image, "fixed_source")
    # Build static ROIs
    # _________________
    roi = call_ipt_func(
        ipt_id="IptRoiManager",
        source=wrapper,
        function_name="generate_roi",
        roi_name='keep_roi',
        left=100,
        width=1885,
        top=300,
        height=1740
    )
    if roi is not None:
        wrapper.add_roi(new_roi=roi)

    roi = call_ipt_func(
        ipt_id="IptRoiManager",
        source=wrapper,
        function_name="generate_roi",
        roi_name='check_roi',
        roi_type='enforce',
        left=840,
        width=400,
        top=1640,
        height=400
    )
    if roi is not None:
        wrapper.add_roi(new_roi=roi)

    roi = call_ipt_func(
        ipt_id="IptRoiManager",
        source=wrapper,
        function_name="generate_roi",
        roi_type='other',
        left=100,
        width=1885,
        top=300,
        height=1740
    )
    if roi is not None:
        wrapper.add_roi(new_roi=roi)

    roi = call_ipt_func(
        ipt_id="IptRoiManager",
        source=wrapper,
        function_name="generate_roi",
        roi_name='open_pot_top',
        roi_type='open',
        left=710,
        width=660,
        top=1990,
        height=50
    )
    if roi is not None:
        wrapper.add_roi(new_roi=roi)

    roi = call_ipt_func(
        ipt_id="IptRoiManager",
        source=wrapper,
        function_name="generate_roi",
        roi_name='safe_roi',
        roi_type='safe',
        left=230,
        width=1550,
        top=350,
        height=1580
    )
    if roi is not None:
        wrapper.add_roi(new_roi=roi)

    # Pre process image (make segmentation easier)
    # ____________________________________________
    wrapper.current_image = call_ipt(
        ipt_id="IptExposureChecker",
        source=wrapper,
        return_type="result",
        overexposed_limit=175,
        over_color='blue_cabin',
        underexposed_limit=35,
        under_color='blue_cabin'
    )

    wrapper.current_image = call_ipt(
        ipt_id="IptPartialPosterizer",
        source=wrapper,
        return_type="result",
        blue_color='blue_cabin',
        post_blue_value=45
    )

    if print_mosaic:
        wrapper.store_image(wrapper.current_image, "pre_processed_image")
    # Build coarse masks
    # __________________
    mask_list = []
    current_mask_ = call_ipt(ipt_id="IptThreshold", source=wrapper, return_type="result", max_t=100)
    mask_list.append(current_mask_)

    current_mask_ = call_ipt(
        ipt_id="IptThreshold", source=wrapper, return_type="result", channel='b', min_t=125
    )
    mask_list.append(current_mask_)

    # Merge masks
    func = getattr(wrapper, "multi_and", None)
    if func:
        wrapper.mask = func([mask for mask in mask_list if mask is not None])
        wrapper.store_image(wrapper.mask, f"mask_multi_and")
        if print_mosaic:
            wrapper.store_image(wrapper.mask, "coarse_mask")
    else:
        wrapper.error_holder.add_error("Unable to merge coarse masks")
        return

    # ROIs to be applied after mask merging
    # _____________________________________
    handled_rois = ['keep', 'delete', 'erode', 'dilate', 'open', 'close']
    rois_list = [
        roi for roi in wrapper.rois_list
        if roi.tag in handled_rois and not (roi.target and roi.target != 'none')
    ]
    wrapper.mask = wrapper.apply_roi_list(img=wrapper.mask, rois=rois_list, print_dbg=True)
    if print_mosaic:
        wrapper.store_image(wrapper.mask, "mask_after_roi")

    # Clean merged mask
    # _________________
    wrapper.mask = call_ipt(
        ipt_id="IptKeepLinkedContours",
        source=wrapper,
        return_type="result",
        tolerance_distance=50,
        tolerance_area=100
    )
    if wrapper.mask is None:
        return

    if print_mosaic:
        wrapper.store_image(wrapper.mask, "clean_mask")

    # Check that the mask is where it belongs
    # _______________________________________
    if print_images:
        res = True
        enforcers_list = wrapper.get_rois({"enforce"})
        for i, enforcer in enumerate(enforcers_list):
            mask = wrapper.mask.copy()
            mask = wrapper.keep_roi(mask, enforcer)
            partial_ok = np.count_nonzero(mask) > 0
            res = partial_ok and res
            if partial_ok:
                roi_img = np.dstack((np.zeros_like(mask), mask, np.zeros_like(mask)))
            else:
                roi_img = np.dstack((np.zeros_like(mask), np.zeros_like(mask), mask))
            background_img = cv2.bitwise_and(wrapper.mask, wrapper.mask, mask=255 - mask)
            img = cv2.bitwise_or(roi_img, np.dstack((background_img, background_img, background_img)))
            enforcer.draw_to(img, line_width=4)
            wrapper.store_image(img, f'enforcer_{i}_{enforcer.name}')
        if not res:
            return
    else:
        enforcers_list = wrapper.get_rois({"enforce"})
        for i, enforcer in enumerate(enforcers_list):
            mask = wrapper.mask.copy()
            mask = wrapper.keep_roi(mask, enforcer)
            if np.count_nonzero(mask) == 0:
                return

    # Extract features
    # ________________
    wrapper.current_image = wrapper.retrieve_stored_image('exposure_fixed')
    wrapper.csv_data_holder = AbstractCsvWriter()
    current_data = call_ipt(ipt_id="IptAnalyseObservation", source=wrapper, return_type="data")
    if isinstance(current_data, dict):
        wrapper.csv_data_holder.data_list.update(current_data)
    else:
        wrapper.error_holder.add_error("Failed to add extracted data")

    current_data = call_ipt(ipt_id="IptAnalyzeChlorophyll", source=wrapper, return_type="data")
    if isinstance(current_data, dict):
        wrapper.csv_data_holder.data_list.update(current_data)
    else:
        wrapper.error_holder.add_error("Failed to add extracted data")

    current_data = call_ipt(ipt_id="IptAnalyzeColor", source=wrapper, return_type="data")
    if isinstance(current_data, dict):
        wrapper.csv_data_holder.data_list.update(current_data)
    else:
        wrapper.error_holder.add_error("Failed to add extracted data")

    current_data = call_ipt(ipt_id="IptAnalyzeObject", source=wrapper, return_type="data")
    if isinstance(current_data, dict):
        wrapper.csv_data_holder.data_list.update(current_data)
    else:
        wrapper.error_holder.add_error("Failed to add extracted data")

    # Save CSV
    if dst_folder and (len(wrapper.csv_data_holder.data_list) > 0):
        with open(
            os.path.join(dst_folder, '', wrapper.file_handler.file_name_no_ext + '.csv'), 'w', newline=''
        ) as csv_file_:
            wr = csv.writer(csv_file_, quoting=csv.QUOTE_NONE)
            wr.writerow(wrapper.csv_data_holder.header_to_list())
            wr.writerow(wrapper.csv_data_holder.data_to_list())

    # Build mosaic
    # ____________
    if print_mosaic:
        wrapper.store_mosaic = 'result'
        wrapper.mosaic_data = np.array([['fixed_source', 'pre_processed_image', 'coarse_mask'],
                                        [
                                            'mask_after_roi', 'clean_mask',
                                            wrapper.draw_image(
                                                src_image=wrapper.current_image,
                                                src_mask=wrapper.mask,
                                                background='bw',
                                                foreground='source',
                                                bck_grd_luma=120,
                                                contour_thickness=6,
                                                hull_thickness=6,
                                                width_thickness=6,
                                                height_thickness=6,
                                                centroid_width=20,
                                                centroid_line_width=8
                                            )
                                        ]])
        wrapper.print_mosaic(padding=(-4, -4, -4, -4))

    print("Done.")


if __name__ == "__main__":
    main()
```
